# Proposing the solar wind-energy-flux hypothesis as a driver of interannual variation in tropical tree reproductive effort tropical tree reproduction

**DOI:** 10.1101/606871

**Authors:** J. Aaron Hogan, Christopher J. Nytch, John E. Bithorn, Jess K. Zimmerman

## Abstract

A growing body of research documents how the El Niño Southern Oscillation (ENSO) results in short-term changes in terrestrial environmental conditions, with the potential to drive ecosystem processes as the duration and severity of ENSO events increases with anthropogenic climate change. An ENSO positive phase results in anomalous patterns of rainfall and temperature throughout the tropics that coincide with leaf flush and increased fruit production in tropical forests worldwide. However, our understanding of possible mechanisms underlying this natural phenomenon is limited. Furthermore, flowering in tropical trees anticipates ENSO development, motivating the continued search for a global phenological cue for tropical angiosperm reproduction. We propose the solar energy flux hypothesis: that a physical energy influx in the Earth’s upper atmosphere and magnetosphere generated by a positive anomaly in the solar wind preceding ENSO development, cues tropical trees to increase allocation of resources to reproduction. We show that from 1994-2013, the solar wind energy flux into the Earth’s magnetosphere (E_in_) is more strongly correlated with the number of trees in fruit or flower in a Puerto Rican wet forest than the Niño 3.4 climate index, despite Niño 3.4 being a previously identified driver of interannual increases in reproduction. We discuss the idea that changes in the global magnetosphere and thermosphere conditions via solar wind-effects on global atmospheric circulation, principally a weaker Walker circulation, cue interannual increases tropical tree reproduction. This may be a mechanism that synchronizes the reproductive output of the tropical trees to changes in environmental conditions that coincide with ENSO. Thus, space weather patterns may help explain terrestrial biological phenomena that occur at quasi-decadal scales.

## Global climate oscillations and variation tropical forest reproduction

Interannual fluctuations in global climate, such as ENSO (McPhaden et al. 2006; Moy et al. 2002; Power et al. 2013; Vecchi et al. 2006), influence tropical forest energy flux (e.g. change in forest temperature or rates of nutrient cycling, Malhi and Wright 2004, Levin et al. 2018) including biomass accretion (Phillips et al. 1998), carbon dynamics (Brienen et al. 2015), and reproductive phenologies of tropical trees (Chang - Yang et al. 2016; Lasky et al. 2016; Pau et al. 2018; Wright and Calder□n 2006; Zimmerman et al. 2007; Zimmerman et al. 2018). As tropical forests account for approximately one-third of the global carbon cycle (Beer et al. 2010, Pan et al 2011), understanding how such interannual climate drivers affect their reproduction is important for long-term projections of tropical forests dynamics, including their carbon storage potential. Changes in the local environment, such as solar irradiance, temperature, and precipitation, only partly describe the variability in leaf and reproductive phenologies of tropical trees (Chapman et al. 2018; Chen et al. 2018; Lasky et al. 2016; Wright and Calderón 2018). A positive ENSO anomaly increases soil moisture deficit, solar radiation, and vapor pressure deficit (Detto et al. 2018; He et al. 2018), during which, tropical trees exhibit increased reproduction (Chang - Yang et al. 2016; Pau et al. 2018; Wright and Calder□n 2006; Zimmerman et al. 2018). For example, the masting of Asian Dipterocarps in ENSO years can increase up to 8-fold from non-ENSO years (Curran and Leighton 2000; Chen et al. 2018).

Yet, how can it be that species mast at the regional and global scales (Koenig and Knops 1998)? How can forests from across the world respond similarly to global-scale interannual climate cycles like ENSO (Asner et al. 2000)? If it were merely a function of resource allocation within trees and response to local abiotic-environmental drivers, one would predict canopy-damaging disturbance, e.g. frost events (Chang - Yang et al. 2016), or hurricanes (Zimmerman et al. 2018), to disrupt the coordination of ENSO and increased reproductive activity among forests. Yet globally, despite disturbance, trees coordinate resource allocation to maximize reproductive effort that coincides almost perfectly with the timing of environmental conditions conducive to high rates of seed survival and germination (i.e. the high light and dry conditions of a positive ENSO). We posit that trees can anticipate ENSO using a yet undetermined physiological cue, which is related to a physical energy increase in the Earth’s magnetosphere and upper atmosphere.

On one hand, tropical trees may be adapted to trade-off the timing of leaf flush and fruiting phenologies to maximize the exploitation of solar insolation by new leaves with the investment of sugars into fruits under dry conditions to avoid drought stress (Detto et al. 2018). A developing ENSO event may trigger a switch in resource investment from leaves to flower and fruits. On the other, by synchronizing an increased volume of fruit production among individuals or species across years, the per-seed cost of negative density-dependent effects, such as exposure to fungal pathogens or seed predation, is minimized (Curran and Leighton 2000; Janzen 1970; Pearse et al. 2016). There is an inherent fitness implication to this adaptive behavior at the population and community scales (Crawley and Long 1995; Kelly et al. 2000), and evolution may have selected for species with higher sensitivity to phenological cues that result in community synchrony in reproduction via increased survival of their seeds. However, in regard to reproductive effort (i.e. timing and output), individual trees are likely insensitive to the benefits of post-fruit production density-dependent effects (Connell and Green 2000; Crawley and Long 1995; Salisbury 1942), and the gestation time for most tropical fruits is several months, suggesting a separate abiotic cue that allows them to forecast the onset of ENSO. We contend that this cue may be electromagnetic or energetic in nature and is likely mediated through subtle changes in the Earth’s upper atmosphere and magnetosphere (i.e. temperature, vapor pressure, atmospheric conductivity) that results from increased energy input from the solar wind.

## The solar wind and the El Niño Southern Oscillation

Recently, using a new modeling approach for solar wind dynamics, He et al. (2018) discovered both a 2-4-year interannual and 11-year quasi-decadal periodicity in E_in_. They further identified a statistically significant relationship between the mean annual strength of the solar wind and subsequent early winter ENSO onset, concluding that increased E_in_ leads to cascading changes to the Earth’s atmospheric and oceanic currents. Such cascading effects include a weakening in the Walker circulation (Rasmusson and Carpenter 1982b, He et al. 2018; Vecchi et al. 2006) and a strengthening in the Bjerknes feedback (Rasmusson and Carpenter 1982b, He et al. 2018; McPhaden et al. 2006), which allow a positive ENSO to develop, however, these climate feedbacks have not, as of yet, been undoubtedly linked to positive anomalies in the solar wind energy flux into the Earth system (Hocke 2009) or to plant lifecycles in any capacity. From 1964 to 2013, positive anomalies in E_in_ preceded the onset of sea level pressure and Walker circulation anomalies by several months to a year (He et al. 2018). Moreover, tropical cyclone and geomagnetic activity have been linked to E_in_ positive anomalies potentially via uneven heating of the thermosphere (80-100 km above sea level) from increased solar wind activity (Li et al 2018). This describes a potential mechanism by which tropical trees may anticipate ENSO and adds to evidence that solar wind-Earth system interactions may potentially drive interannual and quasi-decadal fluctuations in the Earth’s climate, including ENSO (Hocke 2009).

Simple time series correlations between the community phenological response (the number of species in fruit or flower) for two extensively-studied Neotropical forests, Barro Colorado Island (BCI), Panama and Luquillo, Puerto Rico have revealed negative lags, particularly with respect to temperature (Wright and Calder□n 2006; Zimmerman et al. 2007; Zimmerman et al. 2018). Thus, it is reasonable to consider that tropical trees anticipate ENSO irrespective of any cue in the immediate environment (although the cue may be environmentally regulated to some degree through potential positive or negative feedbacks); i.e. to maximize fruit production by the time an ENSO has fully developed, trees must anticipate the event and shift the allocation of resources to reproduction well in advance of local environmental changes. Based on these observations, we propose a new hypothesis that integrates space weather.

## Hypothesis

### Explicitly, the solar wind energy flux hypothesis as a cue for tropical tree reproduction states that

> due to positive energy anomalies in solar wind energy and its interaction with the Earth system’s upper atmosphere (the magnetosphere and thermosphere), tropical trees are physiologically cued to shift resource allocation away from photosynthesis and growth and toward maximum fruit production in preparation for the favorable environmental conditions that will develop during ENSO.

We posit that the physiological mechanism by which tropical trees are cued is linked to changes in tropical atmospheric circulation currents, temperature, vapor pressure which feedback to affect soil moisture (i.e. land-atmospheric coupling) that are possibly driven by solar wind anomalies. Atmospheric conditions and soil moisture are both affected under positive ENSO conditions (Rasmusson & Carpenter 1982a, Sun et al. 2014; Detto et al. 2018, Levin et al. 2018), and have feedbacks on tropical forest productivity (Asner et al. 2000, Levin et al. 2018), therefore it is reasonable to hypothesize they may exert some effect on tropical forest reproduction. Next, we provide a case study that illustrates a stronger correlative relationship to solar-wind energy anomalies than ENSO itself, as preliminary evidence.

## Methods

### Measuring forest phenology using seed traps at Luquillo

Fortnightly surveys of all plant reproductive parts were conducted for 120 stationary traps located in the 16-ha Luquillo Forest Dynamics Plot (18°20’ N, 62°49’ W) in the northwest section of the Luquillo Experimental Forest in eastern Puerto Rico from March 1992 through 2015. The forest community at Luquillo is a Caribbean subtropical montane forest (below 600 m.a.s.l.) dominated by the palm *Prestoea acuminata* var. *montana* (Graham) A.J.Hend. & Galeano and Dacryodes excelsa Vahl. (Burseraceae), with a species richness of 44 tree species per ha ≥1 cm diameter at 1.3 m height (Thompson et al. 2002). Flower presence-absence and seed and fruit abundances are recorded by species for each trap. Fruits were converted to seed abundances using the number of seed per fruit determined for each species (Wright and Calderón 2006; Zimmerman et al. 2007). We limit analyses to species that were recorded in greater than six traps, ensuring that multiple individuals were sampled (Zimmerman et al 2007). Altogether in 23 years of monitoring, 89 species were found in flower and 76 were found in seed or fruit, with 71 species recorded in both flower and fruit (Zimmerman et al. 2018). Seed-trap size was increased from 0.16 to 0.5 m^2^ in 2006, and traps were run concurrently for a year. Seed abundances from the smaller traps were corrected based on the slope of the regression for pairwise flower presences (1.26) and seed abundances (1.61, for additional details see methods in Zimmerman et al. 2018).

### Wavelet analysis

We used a continuous multivariate Morlet wavelet transformation to calculate the time-scale of coherent patterns in community flower and seed abundances. Wavelet analysis allows for the identification of synchronous and compensatory trends at the scale of climate oscillation recurrence (i.e. time scales greater than one year; the scale was specified from 0.5 to 10 years for the analysis). The continuous multivariate wavelet transform is: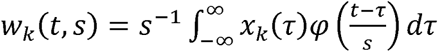, where *φ*(*t*) is the wavelet function and *x*_*k*_(*τ*) is the presence count of flowers or abundance of seeds for the *k*^th^ species at time *τ*, and *s* is wavelet scale. The Morlet wavelet function used is (Morlet et al. 1982): 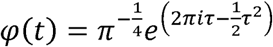. Once data are in wavelet form, one can compute wavelet modulus ratio, a measure of time series coherency at a given time scale, represented by the coefficient of the aggregate temporal variation over the marginal temporal variation of individual species. Coherency via the wavelet modulus ratio was estimated with respect to wavelet scale using 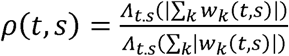, where 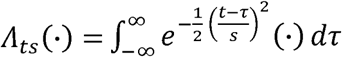 and | · | is the complex norm (i.e. modulus of a complex number) (Keitt 2008; 2014). Wavelet modulus ratios near 1 correspond to community synchrony, while ratios near 0 show community compensation. Statistical significance was determined using a phase-randomization, non-parametric bootstrapping method using 999 randomizations, where the observed wavelet modulus ratio was compared to a null distribution in which wavelet scale and response variables are randomized (wmr.boot function in Keitt 2014). Analyses were carried out in R v.3.2.5 (R Core Team 2016) using the ‘mvcwt’ package (Keitt 2014).

### Seasonal detrending of the phenology data and correlations with solar wind and climate indices

Using the same fortnightly survey data, the phenological response of the forest was defined as the number of species in either fruit of fllower at the monthly-scale, because that is the scale for ENSO climate indices. We seasonally detrended monthly time series of the number of species in flower and seed using seasonal-trend decomposition by loess (Cleveland et al. 1990, see Appendix 2 of Zimmerman et al. 2018). From the trend, annual open, high, low, close (OHLC) moving averages were computed. We obtained annual normalized E_in_ values from 1993-2012 from He et al. (2018). Space weather data on sunspot number (SSN), the solar radio frequency at 10.7 cm (F107), and total solar irradiation (TSI) were downloaded from the NASA ONMIweb database (omniweb.gsfc.nasa.gov) and monthly Niño 3.4 data were obtained from NOAA (esrl.noaa.gov/psd/enso/dashboard.html). SSN, F107, and TSI are commonly used as proxies for solar weather activity, which is primarily a function of the surface mixing of Sun and solar flaring activity. We compared the OHLC moving averages to annualized mean anomalies for E_in_, SSN, F107, TSI, and ENSO 3.4 from 1992-2012. All indices were normalized, and the average annual anomaly was calculated. Pearson correlations between the seasonally-detrended OHLC (open, high, low, close) average of the number of species in flower or seed and the normalized anomaly for the E_in_ and ENSO 3.4 were done using annual data from 1993-2012 (df = 18).

## Results

### Assessing the interannual synchrony in tropical tree reproductive output

To help visualize the community composition of fruits and flowers, and display time-scales at which coherency occurs, we present a Morlet wavelet analysis of flower and fruit abundances. It illustrates supra-annual coherency in a 22-year time series of flower and fruit production at Luquillo (Fig. 1). Community synchrony in flower and seed production occurred at the interannual scales consistent with ENSO. This occurred despite a large disturbance in September 1998, Hurricane Georges which nullified flower and fruit production of the community for roughly three months. More temporally-consistent community synchrony is evident for flowers than for seeds (Fig. 1a) and extended from a timescale of ca. 2 years and beyond. The entire top portion of Fig. 1a is statistically significant as the bold lines delimiting areas of significance using α=0.05 do not connect along the upper boundary. For seed production, the strongest community synchrony occurs between 2 and 5 years and is strongest during the 1997-1998 ENSO (Fig. 1b). Therefore, flower production in the community is regularly synchronized, but seed production is only synchronized interannually and coincides with the timing and periodicity of ENSO.

**Figure 1:**
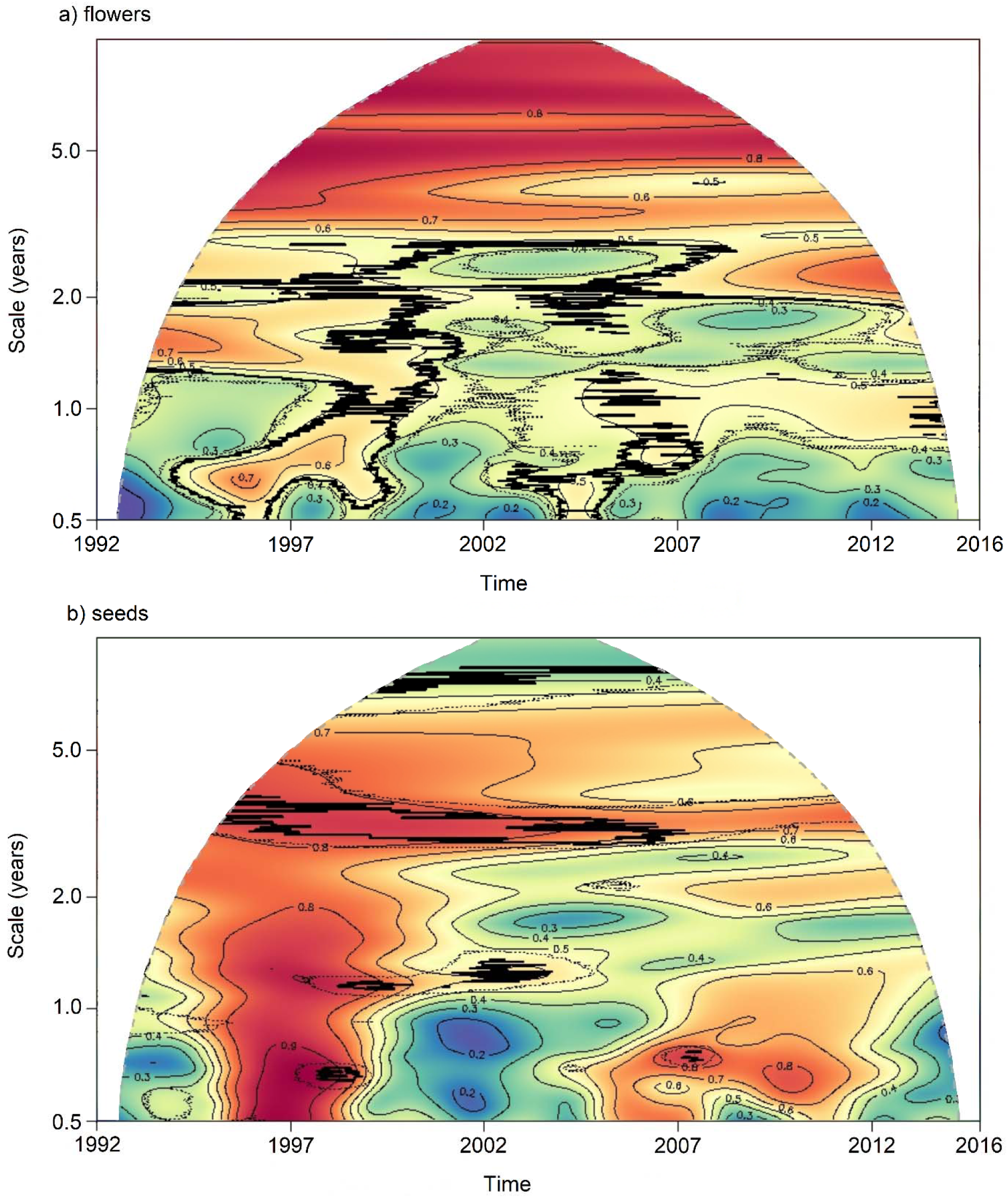
Morlet wavelet modulus ratios over time using seed trap abundances of (a) flowers and (b) seeds for the tree community of the Luquillo Forest Dynamics Plot, Puerto Rico. The vertical axis (log scale) represents the scale of the Morlet wavelet modulus, ranging from a half to 10 years. Colors within the heatmaps denote the magnitude of the wavelet modulus ratio scaled from 0 (dark blue) corresponding to strong compensatory dynamics to 1 (dark red) denoting strong community synchrony. Note that the community synchrony (i.e. the red area) extends down to the smallest scale in Fig. 1b, illustrating the effect of Hurricane Georges in 1998. Contours are wavelet modulus ratio-magnitude isotherms. Dotted and bold lines delimit areas of statistical significance at α = 0.1 and 0.05, respectively using 999 bootstrapped resamples (see Methods).

### Correlation of forest reproduction with the solar wind energy flux

In this context, we were interested in preliminarily exploring how E_in_ related to forest-level flower and seed production. In previous work, ENSO 3.4 was the strongest correlated El Niño-related climate index to the phenology of the trees in Luquillo (Zimmerman et al. 2018). Pearson correlations between the normalized annual E_in_ anomalies and open, high, low, close (OHLC) annual average of the seasonally-detrended number of species in flower (r = −0.635, t = −3.48, df = 18, *p =* 0.0026) and seed (r = 0.443, t = 2.97, df = 18, *p* = 0.0504) were both statistically significant. Those relationships were stronger than their respective correlations with normalized annual anomalies of ENSO 3.4 (flowers: r =0.019, t = 0.08, df = 18 *p* = 0.938, seeds: r = 0.253, t = 1.11, df = 18, *p* = 0.28). Correlations of other measures of space weather activity (SSN. F107, and TSI) with the OHLC annual average of the seasonally-detrended annual number of species in flower or seed were like those of E_in_, but slightly weaker (Fig. 2).

**Figure 2:**
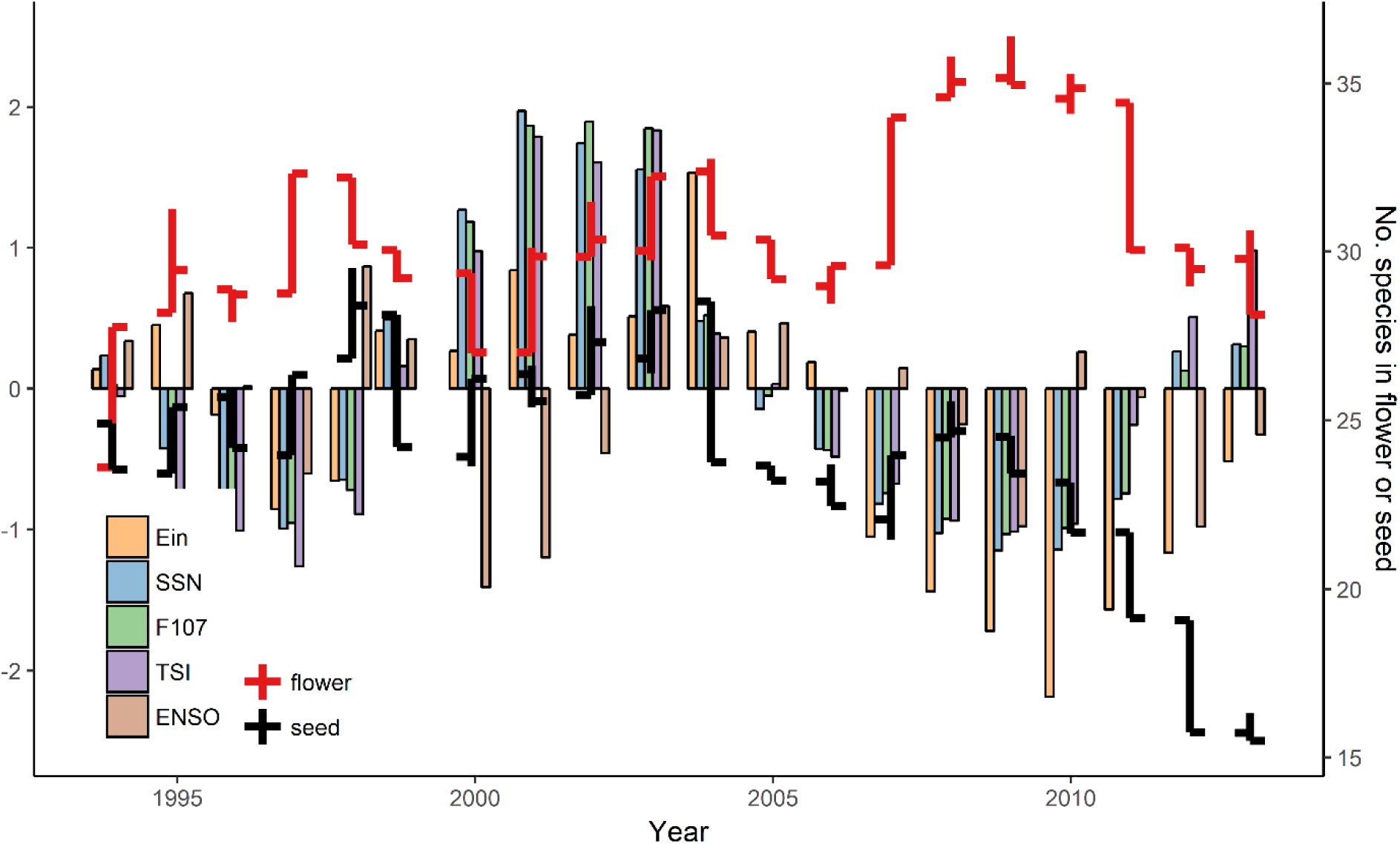
Annual normalized E_in_, SSN, F107, TSI and ENSO 3.4 indices from 1993-2013 with OHLC (Open: left facing horizontal line, High: maximum of annual range, Low: minimum, Close: right facing horizontal line) bars for the seasonally-detrended number of species in flower (red) or seed (blue) at Luquillo, Puerto Rico. Years in which fewer species producing seed coincide with years with negative normalized index values related to solar energy flux into the atmosphere (E_in_, SSN, TSI, and F107) (i.e. less energy input from the solar wind into the Earth system). See supplement for complete lagged time-series correlations.

## Concluding remarks

The community synchrony in the abundance of flowers and fruits counts at 4-6-year timescales is consistent with recurring ENSO positive anomalies (Fig.1). Therefore, the signal of ENSO on community synchrony in the reproductive effort of tropical trees in Luquillo exists despite a dynamic tree community situated in an aseasonal, disturbance-affected forest (Hogan et al. 2016; Hogan et al. 2018; Zimmerman et al. 2007). A positive ENSO has no effect on rainfall at Luquillo but decreases temperature (strongest relationship at 2 to 4-month lead, i.e. before ENSO) and increases solar radiation (photosynthetic photon flux density, strongest relationship at 7-month lag) (Zimmerman et al. 2018). Luquillo is considerably wetter (no month receives <100mm of rainfall) and less seasonal than many tropical forests (e.g., BCI), where ENSO can interact with seasonal dynamics to strengthen or lengthen the dry season (Detto et al. 2018). At the annual scale at Luquillo, relationships with measures of solar wind energy more closely tracked the number of species in flower or seed over time than the ENSO 3.4 anomalies (Fig. 2). E_in_ was found to have no statistically significant relationship with either annual total rainfall or average temperatures (minimum or maximum) yet was strongly related to flower and seed reproductive effort of trees, potentially supporting the idea that changes in global atmospheric circulations relate to interannual variation in reproductive effort of tropical forests.

In conclusion, the roughly 1-year lagged E_in_-ENSO teleconnection occurs most strongly over the eastern Indo-Pacific, creating strongly anomalous Walker circulation diverge there and several convergence centers over the Northeastern Pacific (Rasmussen and Carpenter 1982b, He et al. 2018). These global-scale changes in atmospheric conditions that extend at least 1000 km above the Earth’s surface (He et al. 2018), can be linked to climate, geomagnetic energy and global cyclonic activity (Li et al. 2018). ENSO acts as an established global climatic driver of supra-annual cycles in tropical tree phenology (Chang - Yang et al. 2016; Pau et al. 2018; Wright and Calder□n 2006; Zimmerman et al. 2018, Detto et al. 2018). The mechanism behind the interannual increase in reproductive effort of tropical trees is difficult to identify, owing to the subtlety and complexity of climatic teleconnections, and the need for long-term records of forest reproduction (Abernethy et al. 2018; Pearse et al. 2016; Wright and Calderón 2018). At Luquillo after accounting for seasonal variation, a positive E_in_ anomaly results in fewer species producing flowers, but an increase in species producing seed (Fig. 2). At Luquillo, these patterns appear to be influenced by negative trends in solar energy flux and declines in seeds production after 2005.

Whether this relationship holds at other Neotropical sites remains to be investigated. At BCI, Panama, similar coherency in community reproduction and leaf flush have been observed at 4-7-year timescales (Detto et al. 2018), pointing to the widespread effect of ENSO on tropical tree reproductive effort. However, in contrast to Luquillo, there have been steady increases in fruit and flower production over the 28-year record at BCI (Pau et al. 2018). Notably, we identified no effects between E_in_ and temperature or precipitation for Luquillo using annual data, although previous work has shown ENSO to affect temperature and solar radiation at the sub-annual (i.e. monthly) scale. Likely the interactions between fluctuations in energy into the Earth system from the solar wind and tropical tree reproduction are complex and occur at timescales finer than the annual scale as the solar wind intensity fluctuates with solar activity. Thus, finer temporal resolution in the analysis may prove to be insightful. Furthermore, individual species probably vary in their sensitivity to solar wind energy flux. Future research should work to identify how solar wind energy flux and other measures of solar weather influence tropical tree phenology at broader spatial and temporal scales.

## Supporting information

See supplement for complete lagged time-series correlations.

## Author contributions

J.A.H conceived the idea, analyzed the data, and wrote the manuscript. J.K.Z. and J.E.B collected the data. All authors contributed intellectually and provided insight and comments on manuscript drafts.

## Materials & Correspondence

Materials and correspondence should be addressed to J.A.H.

## Content type

Letter

## Acknowledgments

We thank Drs. Shengping He, and Hui Li from the Key Lab for Space Weather at the Chinese Academy of Sciences for sharing the E_in_ data and providing insights.

## Data availability

Data on the phenology of trees and shrubs from the Luquillo Forest Dynamics Plot can be found on the Luquillo LTER website: dataset 88: http://luq.lter.network/data/luqmetadata88

