## Supplementary material for "Proposing the solar wind-energy-flux hypothesis as a driver of interannual variation in tropical tree reproductive effort tropical tree reproduction": See supplement for complete lagged time-series correlations.

### LUQ_solar wind correlations

*JAH*

*September 13, 2018*

#### LUQ SOLAR WIND CORRELATIONS

```
### LOAD IN THE DATA - fmon and smon timeseries (flowers and seeds byu month)
load("C:/Users/hogie/Dropbox (Personal)/Phenology/LUQ_smon.Rdata")
load("C:/Users/hogie/Dropbox (Personal)/Phenology/LUQ_fmon.Rdata")

library(psych)
library(readr)
solar_wind <- read_table("C:/Users/hogie/Dropbox (Personal)/PHENO_Solar Wind Energy Hyp/Data/Solar Wind
```

```
## Warning: The following named parsers don't match the column names: X6
```

```
### make time series
######### SSN - sun spot number
SSN.ts <- ts(solar_wind[13:276,]$SSN, start = c(1993,1), end = c(2014,12), frequency = 12)
######### F107 - the solar radio flux at 10.7 cm wavelength
F107.ts <- ts(solar_wind[13:276,]$F107, start = c(1993,1), end = c(2014,12), frequency = 12)
### Ein - the solar wind energy flux into the magnetosphere
Ein.ts <- ts(solar_wind[13:276,]$Ein, start = c(1993,1), end = c(2014,12), frequency = 12)


f.mon.ts <- ts(f.mon,  start = c(1993,1), end = c(2014,12), frequency = 12)
s.mon.ts <- ts(s.mon,  start = c(1993,1), end = c(2014,12), frequency = 12)
```

```
xxx1 <- ccf(SSN.ts, f.mon.ts)
```

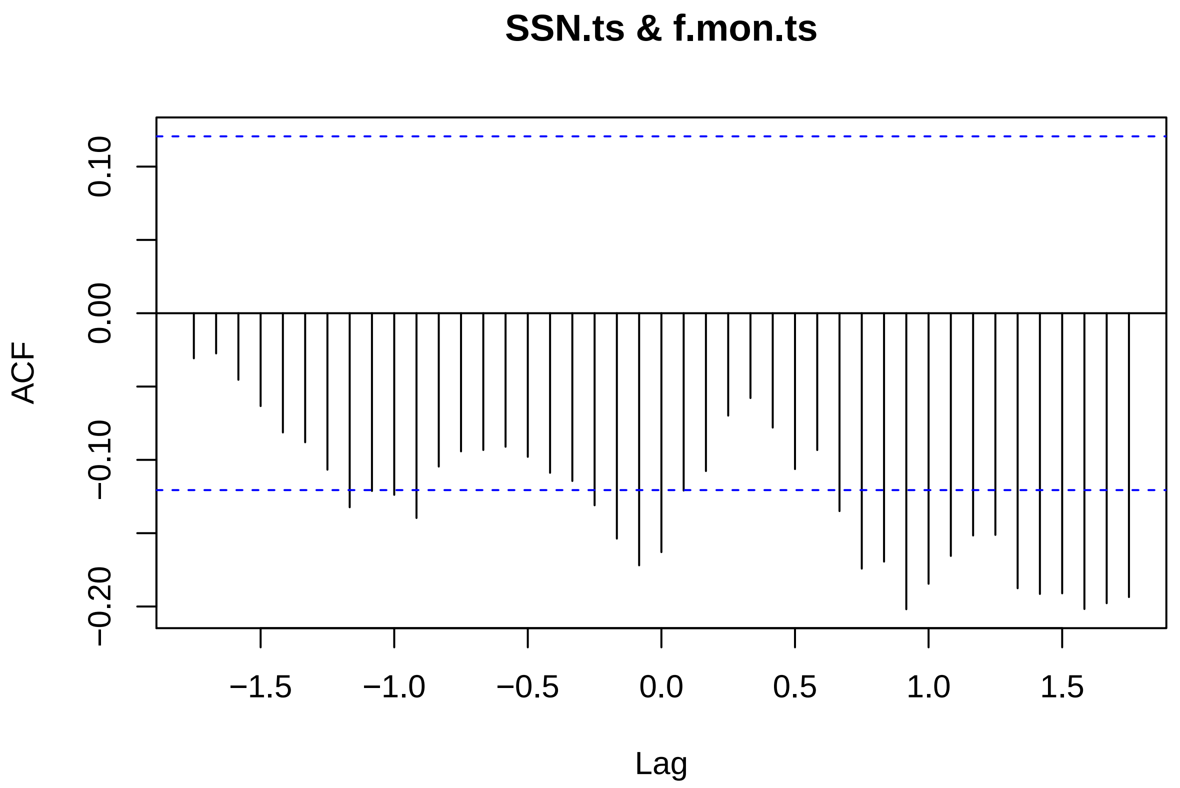


```r
xxx1
```

```
##
## Autocorrelations of series 'X', by lag
##
## -1.7500 -1.6667 -1.5833 -1.5000 -1.4167 -1.3333 -1.2500 -1.1667 -1.0833
##  -0.031  -0.027  -0.045  -0.063  -0.081  -0.088  -0.107  -0.132  -0.121
## -1.0000 -0.9167 -0.8333 -0.7500 -0.6667 -0.5833 -0.5000 -0.4167 -0.3333
##  -0.124  -0.140  -0.105  -0.094  -0.093  -0.091  -0.098  -0.109  -0.114
## -0.2500 -0.1667 -0.0833  0.0000  0.0833  0.1667  0.2500  0.3333  0.4167
##  -0.131  -0.154  -0.172  -0.163  -0.121  -0.108  -0.070  -0.058  -0.078
##  0.5000  0.5833  0.6667  0.7500  0.8333  0.9167  1.0000  1.0833  1.1667
##  -0.106  -0.093  -0.135  -0.174  -0.169  -0.202  -0.185  -0.166  -0.152
##  1.2500  1.3333  1.4167  1.5000  1.5833  1.6667  1.7500
##  -0.151  -0.188  -0.191  -0.191  -0.202  -0.198  -0.194
```

```r
r.test(252, -0.202) #lag 11 months
```

```
## Correlation tests
## Call:r.test(n = 252, r12 = -0.202)
## Test of significance of a  correlation
##  t value -3.26    with probability < 0.0013
##  and confidence interval  -0.32 -0.08
```

```r
xxx2 <-ccf(F107.ts, f.mon.ts)
```

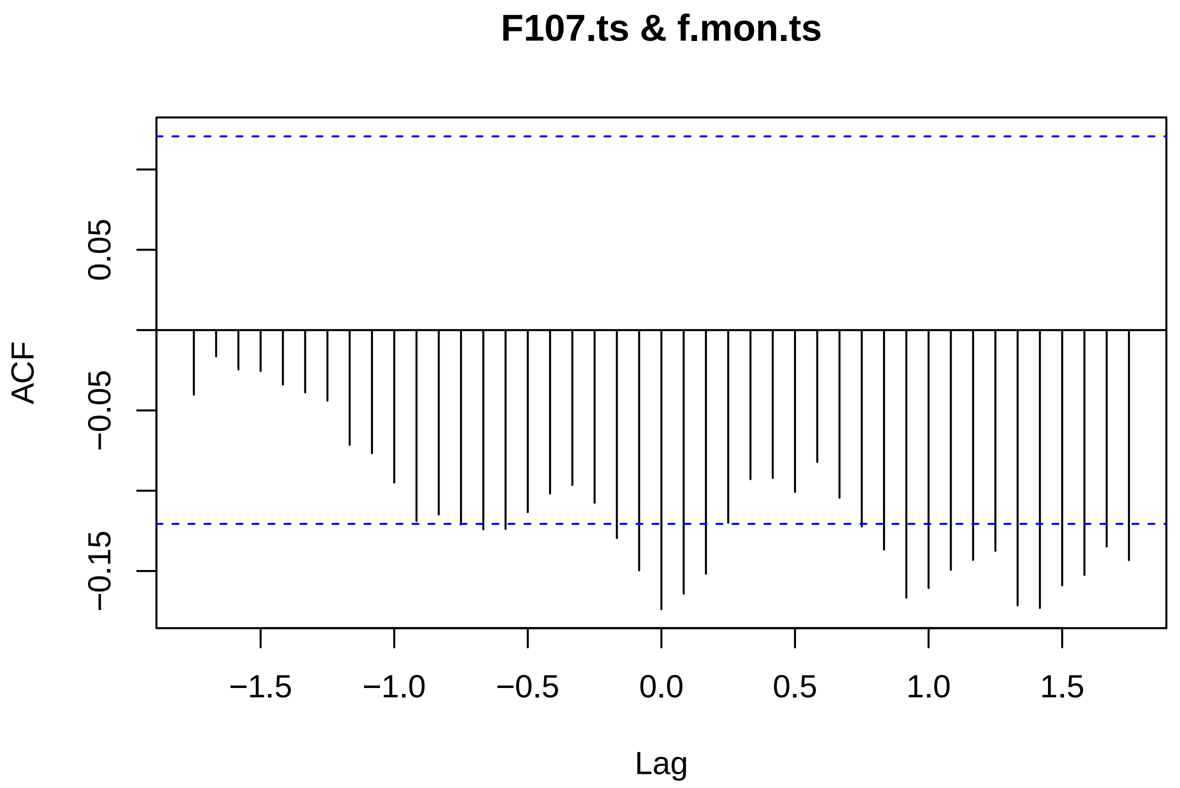


```
xxx2
```

```
##
## Autocorrelations of series 'X', by lag
##
## -1.7500 -1.6667 -1.5833 -1.5000 -1.4167 -1.3333 -1.2500 -1.1667 -1.0833
##  -0.040  -0.016  -0.024  -0.025  -0.034  -0.039  -0.044  -0.071  -0.077
## -1.0000 -0.9167 -0.8333 -0.7500 -0.6667 -0.5833 -0.5000 -0.4167 -0.3333
##  -0.095  -0.119  -0.115  -0.121  -0.124  -0.124  -0.113  -0.102  -0.096
## -0.2500 -0.1667 -0.0833  0.0000  0.0833  0.1667  0.2500  0.3333  0.4167
##  -0.107  -0.130  -0.150  -0.174  -0.164  -0.152  -0.120  -0.093  -0.092
##  0.5000  0.5833  0.6667  0.7500  0.8333  0.9167  1.0000  1.0833  1.1667
##  -0.101  -0.082  -0.104  -0.122  -0.137  -0.167  -0.161  -0.149  -0.143
##  1.2500  1.3333  1.4167  1.5000  1.5833  1.6667  1.7500
##  -0.137  -0.171  -0.173  -0.159  -0.152  -0.135  -0.143
```

```
r.test(252, -0.174) # lag 0 months
```

```
## Correlation tests
## Call:r.test(n = 252, r12 = -0.174)
## Test of significance of a  correlation
##  t value -2.79    with probability < 0.0056
##  and confidence interval  -0.29 -0.05
```

```
xxx3 <- ccf(Ein.ts, f.mon.ts)
```

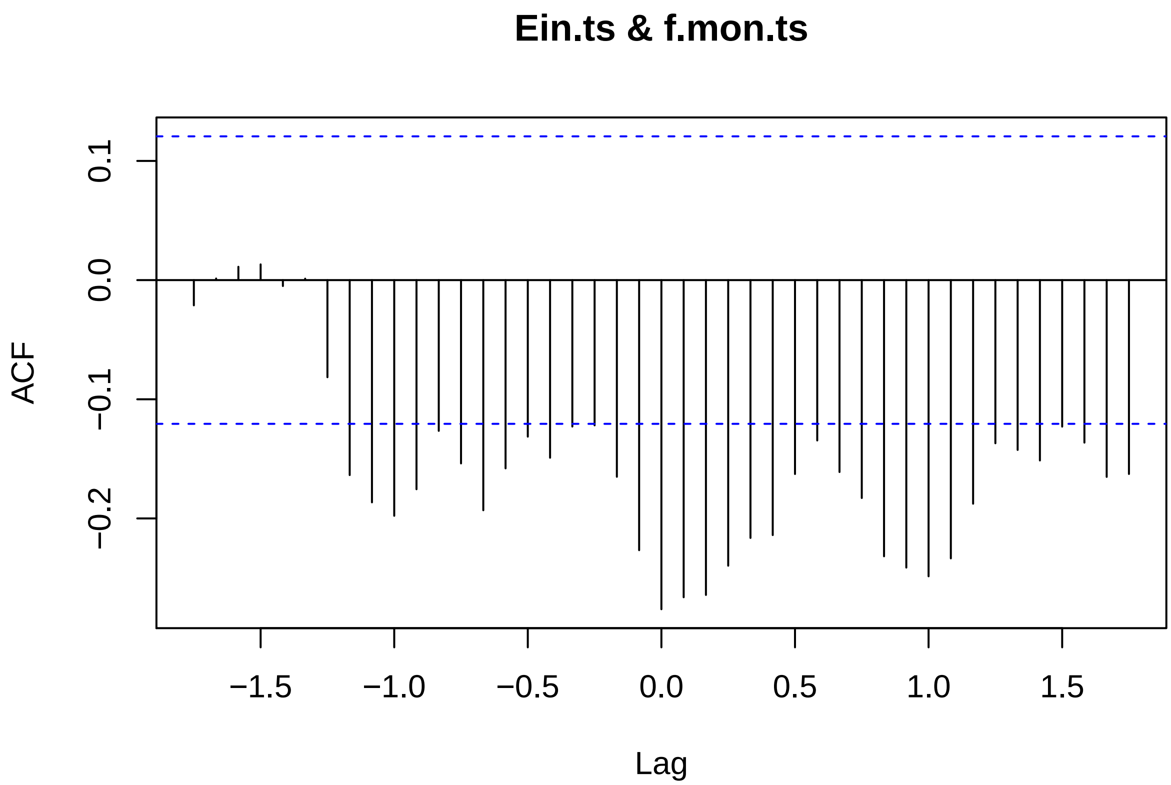


```
xxx3
```

```
##
## Autocorrelations of series 'X', by lag
##
## -1.7500 -1.6667 -1.5833 -1.5000 -1.4167 -1.3333 -1.2500 -1.1667 -1.0833
##  -0.021   0.001   0.011   0.013  -0.005   0.001  -0.081  -0.164  -0.187
## -1.0000 -0.9167 -0.8333 -0.7500 -0.6667 -0.5833 -0.5000 -0.4167 -0.3333
##  -0.198  -0.176  -0.127  -0.154  -0.193  -0.158  -0.131  -0.149  -0.123
## -0.2500 -0.1667 -0.0833  0.0000  0.0833  0.1667  0.2500  0.3333  0.4167
##  -0.122  -0.165  -0.227  -0.276  -0.266  -0.264  -0.240  -0.216  -0.214
##  0.5000  0.5833  0.6667  0.7500  0.8333  0.9167  1.0000  1.0833  1.1667
##  -0.163  -0.135  -0.161  -0.183  -0.232  -0.241  -0.249  -0.234  -0.188
##  1.2500  1.3333  1.4167  1.5000  1.5833  1.6667  1.7500
##  -0.137  -0.143  -0.151  -0.123  -0.136  -0.165  -0.163
```

```
r.test(252, -0.276)
```

```
## Correlation tests
## Call:r.test(n = 252, r12 = -0.276)
## Test of significance of a  correlation
##  t value -4.54    with probability < 8.7e-06
##  and confidence interval  -0.39 -0.16
```

```
xxx4 <- ccf(SSN.ts, s.mon.ts)
```

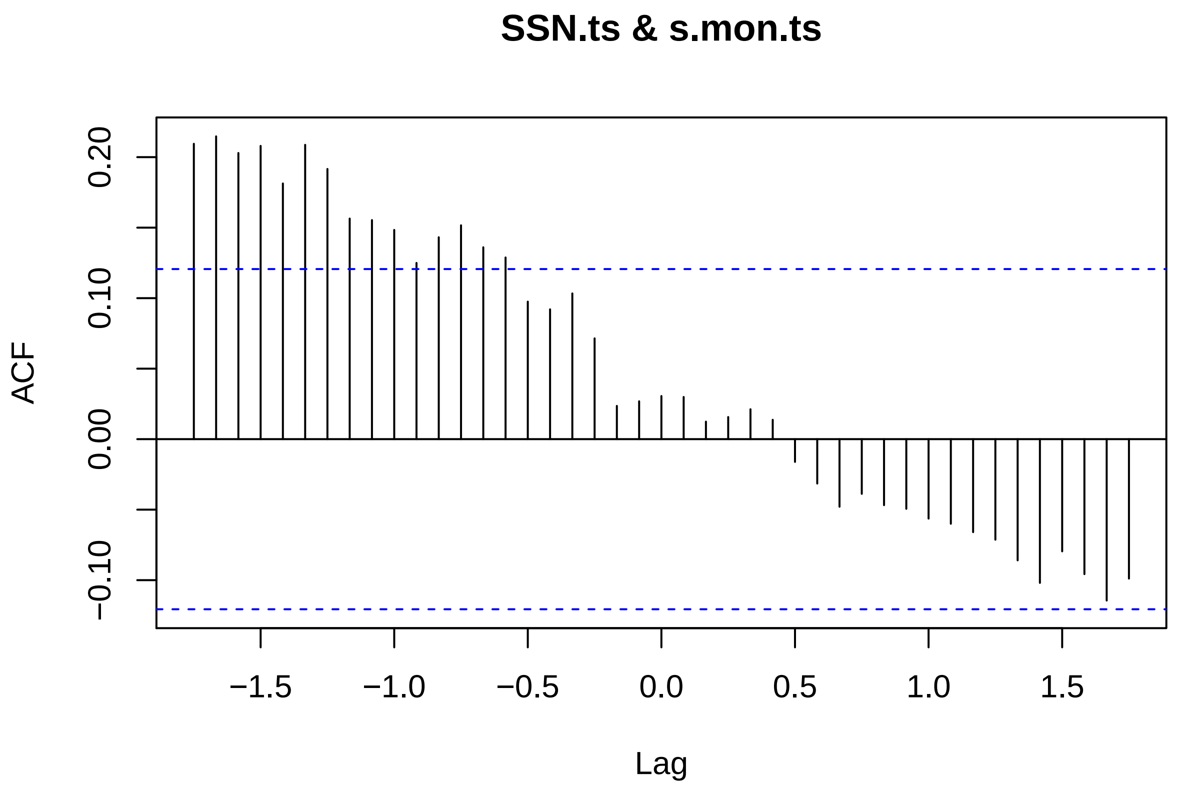


```
xxx4
```

```
##
## Autocorrelations of series 'X', by lag
##
## -1.7500 -1.6667 -1.5833 -1.5000 -1.4167 -1.3333 -1.2500 -1.1667 -1.0833
##   0.209   0.215   0.203   0.208   0.181   0.209   0.192   0.156   0.155
## -1.0000 -0.9167 -0.8333 -0.7500 -0.6667 -0.5833 -0.5000 -0.4167 -0.3333
##   0.148   0.125   0.143   0.152   0.136   0.129   0.098   0.092   0.103
## -0.2500 -0.1667 -0.0833  0.0000  0.0833  0.1667  0.2500  0.3333  0.4167
##   0.071   0.024   0.027   0.031   0.030   0.012   0.016   0.021   0.014
##  0.5000  0.5833  0.6667  0.7500  0.8333  0.9167  1.0000  1.0833  1.1667
##  -0.016  -0.031  -0.048  -0.039  -0.047  -0.049  -0.056  -0.060  -0.066
##  1.2500  1.3333  1.4167  1.5000  1.5833  1.6667  1.7500
##  -0.071  -0.086  -0.102  -0.080  -0.096  -0.114  -0.099
```

```
r.test(252, 0.215)
```

```
## Correlation tests
## Call:r.test(n = 252, r12 = 0.215)
## Test of significance of a  correlation
##  t value 3.48    with probability < 0.00059
##  and confidence interval  0.09 0.33
```

```
xxx5 <- ccf(F107.ts, s.mon.ts)
```

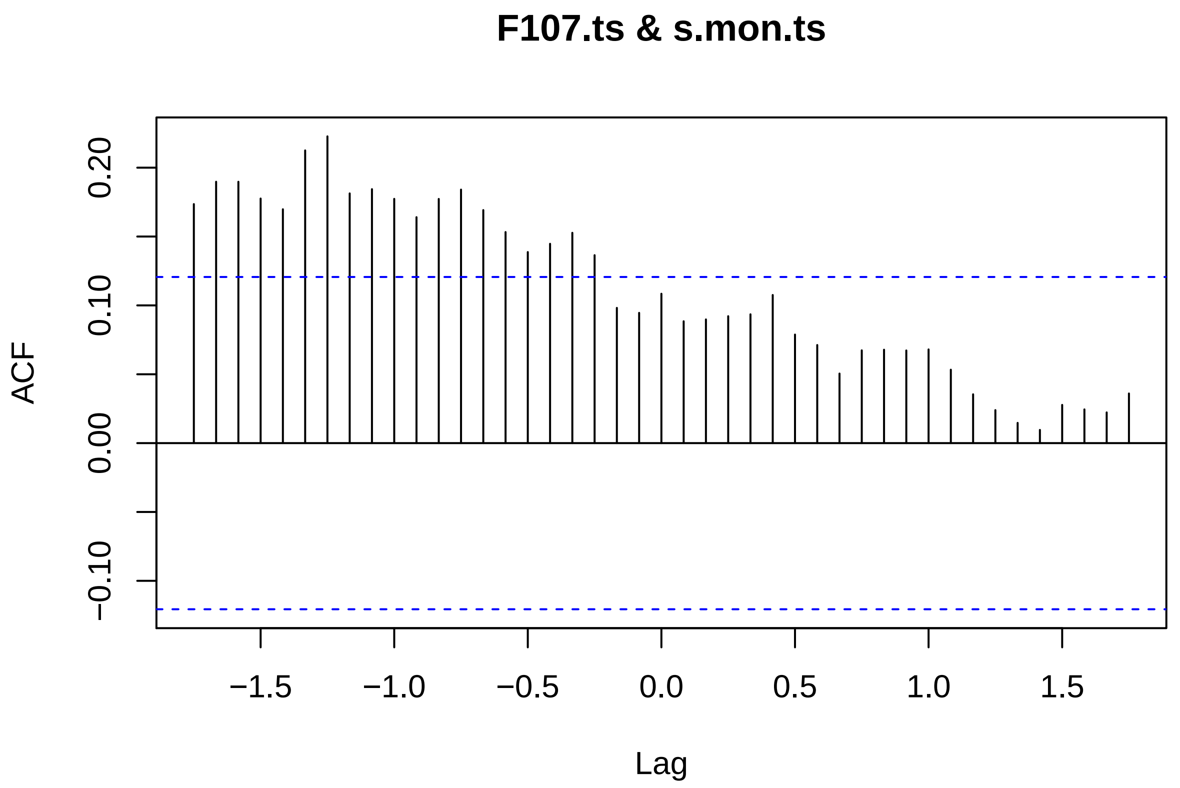


```
xxx5
```

```
##
## Autocorrelations of series 'X', by lag
##
## -1.7500 -1.6667 -1.5833 -1.5000 -1.4167 -1.3333 -1.2500 -1.1667 -1.0833
##   0.174   0.190   0.190   0.178   0.170   0.213   0.223   0.181   0.184
## -1.0000 -0.9167 -0.8333 -0.7500 -0.6667 -0.5833 -0.5000 -0.4167 -0.3333
##   0.177   0.164   0.177   0.184   0.169   0.153   0.139   0.145   0.153
## -0.2500 -0.1667 -0.0833  0.0000  0.0833  0.1667  0.2500  0.3333  0.4167
##   0.136   0.098   0.095   0.108   0.088   0.090   0.092   0.094   0.108
##  0.5000  0.5833  0.6667  0.7500  0.8333  0.9167  1.0000  1.0833  1.1667
##   0.079   0.071   0.051   0.067   0.068   0.067   0.068   0.053   0.035
##  1.2500  1.3333  1.4167  1.5000  1.5833  1.6667  1.7500
##   0.024   0.015   0.010   0.028   0.024   0.022   0.036
```

```
r.test(252, 0.223)
```

```
## Correlation tests
## Call:r.test(n = 252, r12 = 0.223)
## Test of significance of a  correlation
##  t value 3.62    with probability < 0.00036
##  and confidence interval  0.1 0.34
```

```
xxx6 <- ccf(Ein.ts, s.mon.ts)
```

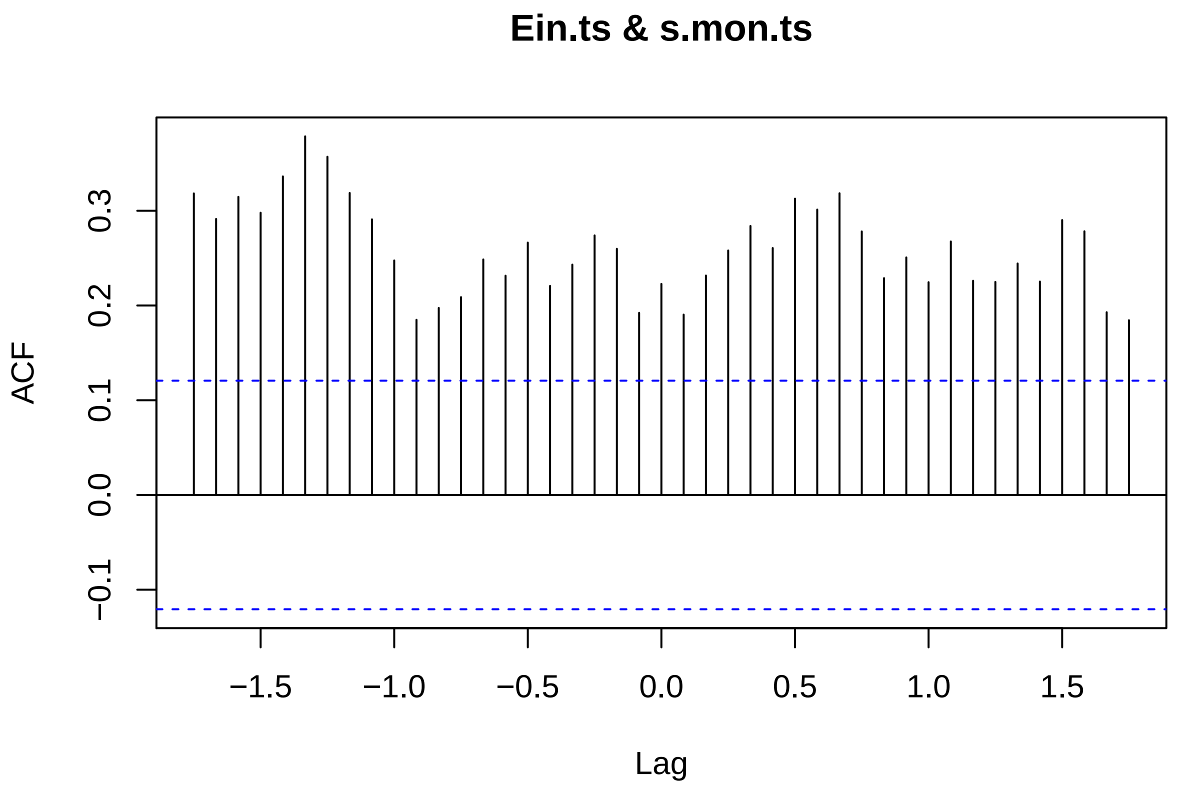


```
xxx6
```

```
##
## Autocorrelations of series 'X', by lag
##
## -1.7500 -1.6667 -1.5833 -1.5000 -1.4167 -1.3333 -1.2500 -1.1667 -1.0833
##   0.318   0.291   0.315   0.298   0.336   0.379   0.357   0.319   0.291
## -1.0000 -0.9167 -0.8333 -0.7500 -0.6667 -0.5833 -0.5000 -0.4167 -0.3333
##   0.248   0.185   0.197   0.209   0.249   0.231   0.266   0.221   0.243
## -0.2500 -0.1667 -0.0833  0.0000  0.0833  0.1667  0.2500  0.3333  0.4167
##   0.274   0.260   0.192   0.223   0.190   0.232   0.258   0.284   0.261
##  0.5000  0.5833  0.6667  0.7500  0.8333  0.9167  1.0000  1.0833  1.1667
##   0.313   0.301   0.318   0.278   0.229   0.251   0.225   0.268   0.226
##  1.2500  1.3333  1.4167  1.5000  1.5833  1.6667  1.7500
##   0.225   0.244   0.225   0.290   0.278   0.193   0.184
```

```
r.test(252, -0.202)
```

```
## Correlation tests
## Call:r.test(n = 252, r12 = -0.202)
## Test of significance of a  correlation
##  t value -3.26    with probability < 0.0013
##  and confidence interval  -0.32 -0.08
```
